# Probabilistic inference of epigenetic age acceleration from cellular dynamics

**DOI:** 10.1101/2023.03.01.530570

**Authors:** Jan. K. Dabrowski, Emma. J. Yang, Samuel. J. C. Crofts, Robert. F. Hillary, Daniel. J. Simpson, Daniel. L. Mccartney, Riccardo. E. Marioni, Eric Latorre-Crespo, Tamir Chandra

## Abstract

The emergence of epigenetic predictors was a pivotal moment in geroscience, propelling the measurement and concept of biological ageing into a quantitative era. However, while current epigenetic clocks have shown strong predictive power, they do not reflect the underlying biological mechanisms driving methylation changes with age. Consequently, biological interpretation of their estimates is limited. Furthermore, our findings suggest that clocks trained on chronological age are confounded by non-age-related phenomena.

To address these limitations, we developed a probabilistic model that describes methylation transitions at the cellular level. Our approach reveals two measurable components, acceleration and bias, that directly relate to perturbations of the underlying cellular dynamics. Acceleration is the proportional increase in the speed of methylation transitions across CpG sites, whereas bias is the degree of global change in methylation affecting all CpG sites uniformly. Using data from 7,028 participants from the Generation Scotland study, we found the age acceleration parameter to be associated with physiological traits known to impact healthy ageing. Furthermore, a genome-wide association study of age acceleration identified four genomic loci previously linked with ageing.

## Introduction

The role of age as the predominant risk factor for cancer, neurodegenerative disease, and cardiovascular disease has motivated research into its underlying cellular mechanisms. Until recently, a major challenge in this field was the lack of a reliable method for accurately measuring biological age. A tipping point was the development of the first comprehensive epigenetic age predictors ^1–4^. These models quantified biological age based on the presence of age-related changes in the DNA methylome of individuals. This development naturally led to the concept of age acceleration, which is commonly defined, for an individual within a cohort, as the residual from the epigenetic clock’s predicted age and their chronological age ^1,5^. The use of epigenetic clocks to measure the rate of biological ageing is now widely employed as they have been shown to capture the impact of various diseases and environmental factors in a single metric ^5,6^.

In recent years, advances in the field have led to numerous improvements. First, population-based cohorts used for training have increased from modest sizes to include and combine large cross-sectional studies with thousands of participants ^3,4,7,8^. Second, the complexity of models has advanced in parallel on two fronts. Machine learning techniques are now used increasingly to develop epigenetic clocks that capture non-linear and interaction dynamics ^9,10^. Furthermore, “composite” or “second generation” clocks have been directly trained on measures of health and longevity. As a result, these clocks have increased associations with a number of diseases as well as overall mortality ^11–13^.

While more sophisticated algorithms and larger cohort sizes have improved the accuracy of epigenetic clocks in predicting chronological age, they do so at the cost of not fully capturing biological information (Zhang et al. 2019). On the other hand, integration of physiological parameters, such as inflammatory markers, into epigenetic clocks shifted their focus from epigenetic ageing toward the prediction of age-related diseases.

Lastly, a common criticism of most commonly applied epigenetic clocks is that the statistical learning approaches used to predict methylation patterns do not necessarily reflect underlying molecular processes. This limitation complicates the interpretation of their biological underpinnings ^6,14^.

In this study, we systematically identify and evaluate design issues with current epigenetic predictors of chronological age. We show that the amount of biological variability captured by these models decreases as the size of cohorts or the complexity of algorithms increases. Further, we highlight how confounding epigenetic processes, such as global change in methylation, can substantially bias predictions of age acceleration.

To overcome these technical limitations and the lack of biological interpretation, we developed a new model of epigenetic age acceleration based on a mathematical representation of the cellular dynamics of methylation change ^15,16^. This model allows mechanistic interpretation of methylation change at CpG sites. Its application predicts two distinct processes that modify the natural progression of epigenetic change over the human lifespan: age acceleration and bias (global change in methylation levels). We provide an efficient method to infer these parameters for each individual using blood-based methylome-wide array data from the Generation Scotland cohort and develop a novel batch correction algorithm to enhance transferability of our model to other cohorts.

We also describe associations between our acceleration parameter and lifestyle factors, prevalent disease outcomes, and risk of all-cause mortality. Further, a genome-wide association study revealed four genomic regions significantly associated with age acceleration.

## Results

### Limitations of clocks trained on chronological age

We first sought to understand why CpGs that do not correlate with chronological age are included in epigenetic age predictors ^14^. Our findings revealed that inclusion of non-age-correlated CpGs (naCpGs, R^2^<0.1) improves chronological age prediction at the expense of capturing variability from disease-related lifestyle factors, as we demonstrate below.

First, we showed the extent to which naCpGs are incorporated in commonly used clocks trained on chronological age (Fig. 1a). We excluded composite epigenetic clocks from our analysis because they are expected to include naCpGs that predict phenotypes other than chronological age ^11,12^. Note that the high number of naCpG sites present in the multi-tissue predictors - Horvath ^1^ and Skin and Blood ^5^ - might be explained in part by the presence of tissue-specific sites. Second, we plotted the age association of all clock CpGs against their association with smoking, which has widespread associations with blood-based DNAm ^17,18^ (Fig. 1b). This revealed naCpGs with methylation levels associated with smoking being included in several clocks. In Fig. 1c, we showed an example of a smoking-correlated CpG site included in the Zhang *et al* clock ^7^. Similarly, in Fig. 1d, we showed a site included in the DeepMAge clock that is highly correlated with alcohol consumption, a correlate of epigenetic age predictors ^9,17^. As DeepMAge is a non-linear neural network predictor, this suggests that incorporation of naCpGs persists even when more complex algorithms are applied.

**Figure 1.**
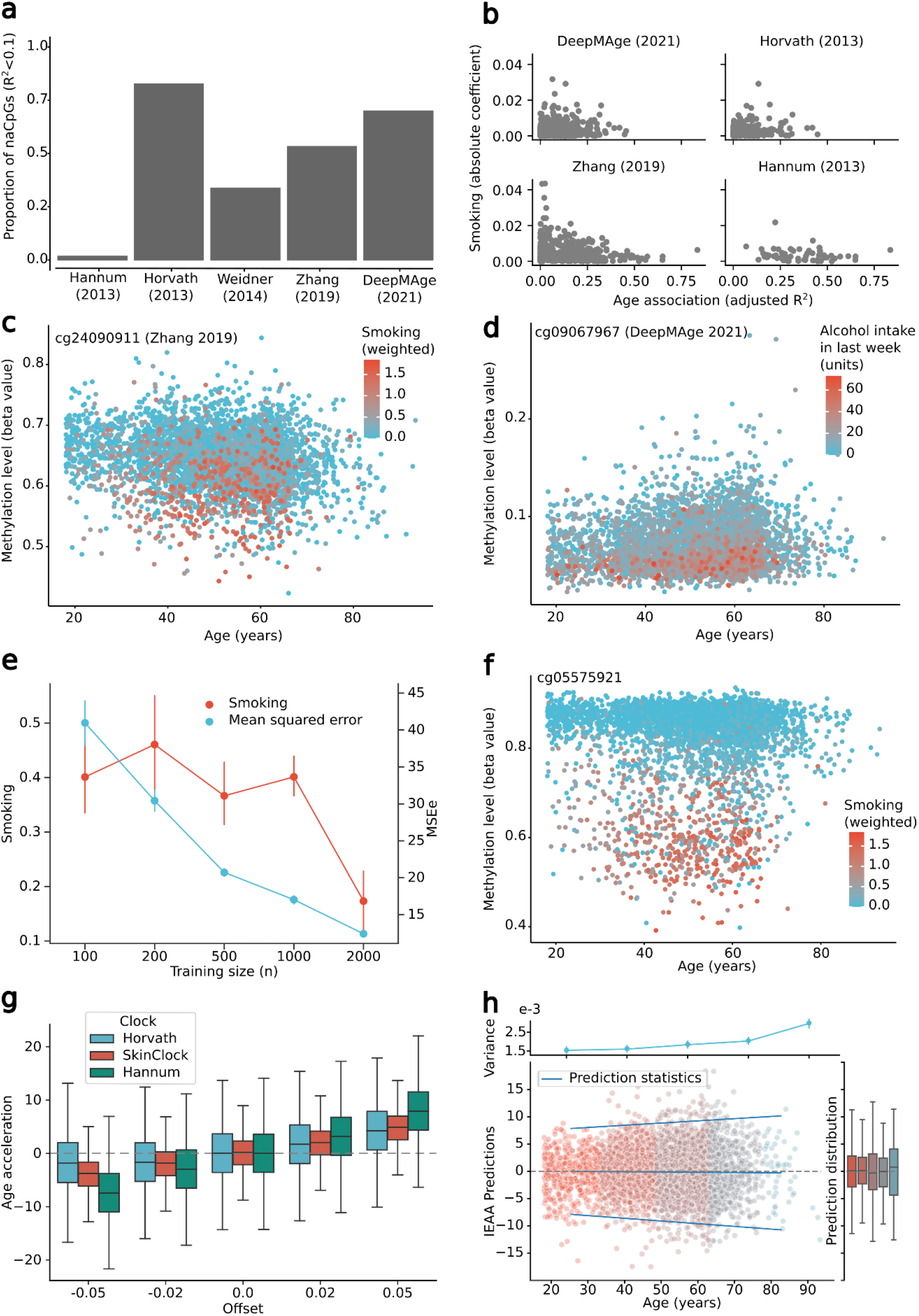
Limitations of current epigenetic predictors. a. Proportion of CpGs included in various epigenetic clocks that are not associated with age. For each CpG site included in these epigenetic clocks, we fitted linear regression models of the form: methylation ~ age. naCPGs were defined as sites below an age-association threshold of R^2^=0.1. b. Comparison between the association of methylation levels with smoking and age on each CpG site included in various epigenetic clocks. Each point corresponds to a site included in a clock. Age association displayed as adjusted R^2^ from linear regressions for each CpG of the form: methylation ~ age. Smoking association shown as the absolute value of the coefficient of smoking (dichotomised as a weighted smoking value of greater than 0.25) from linear regressions for each CpG of the form: methylation ~ smoking + age + sex. Weighted smoking defined in Methods. c. Methylation beta values vs age for a single CpG that is included in the epigenetic clock of Zhang et al. ^7^. Each point represents an individual and is coloured by their level of smoking (defined in Methods). d. Methylation beta values vs age for a single CpG that is included in the DeepMAge clock. Each point represents an individual and is coloured by their level of alcohol intake in the last week. e. Acceleration obtained from bootstrapped lasso linear regressions trained on chronological age for a range of training cohort sizes. Training cohorts were randomly sampled from the GS_set1_ dataset. The results were computed on random test sets (n=2000). The red line, associated to the left y-axis, shows the average, and 95% confidence interval, association between the predicted age accelerations with smoking, given training cohort size. The blue line, associated to the right y-axis, shows the average, and 95% confidence interval, mean squared error in the prediction of chronological age. f. Methylation beta values vs age for a single CpG that is included in all lasso models of chronological age trained with 2000 individuals. Each point represents an individual and is coloured by their level of smoking (defined in Methods). g. Impact of global offsets on the inferred accelerations using Horvath, Skin and Blood and Hannum clocks. Global offsets in methylation were increasingly applied to all sites in GS_set1_ and accelerations were computed as the residual from the predicted age to the chronological age of individuals. Further, accelerations have been shifted for each clock so that the predictions with zero offset are centred at 0. Boxplots show the median and exclusive interquartile range h. Effect of the increase in variance of methylation levels as a function of time in epigenetic predictors. Each point corresponds to the acceleration predicted by Horvath’s clock for every individual in the GS_set1_ cohort. In blue, we show a linear regression of the mean and 2 standard deviations of the predicted accelerations, computed using age bins reflected by the change in colour of points. The right marginal box plot shows the median and exclusive interquartile range of the predicted accelerations by bin. The top marginal plot shows the evolution of the variance across the highest 1000 sites correlating with age in GS_set1_.

To understand why naCpGs are incorporated in linear predictors of chronological age, we investigated how an increase in cohort size impacts age prediction accuracy and association with tobacco smoking (Fig. 1e-f). We used a subset of the Generation Scotland cohort (GS_set1_; n=4450 unrelated individuals with blood-based Illumina EPIC array DNAm) and bootstrapping techniques to train LASSO regression models on training sets of increasing sizes (see Methods). We found that an increased size of the training cohort results in more accurate chronological age prediction, as previously reported (Fig. 1e) ^7^. In contrast, the strength of association between smoking and age acceleration estimated by the bootstrapped models showed a sharp decline for training sets exceeding 1000 individuals. This drop in association correlates with an increasing fraction of naCpG sites being considered, some of which correlate strongly with smoking (Extended Data Fig. 1a). In Fig. 1f, we showed a naCpG strongly associated with smoking (R^2^=0.59, p<0.001) that becomes incorporated in all models trained with 2000 individuals. Next we tested the effect of increasing the prevalence of smokers in training cohorts of fixed size (n=700, see Methods). This test produced analogous results: the strength of association between smoking and age acceleration decreases as the prevalence of smokers increases (Extended Data Fig. 1b-c). Similarly, the naCpG illustrated in Fig. 1f is incorporated in every clock trained with more than 30% of smokers.

**Figure 2.**
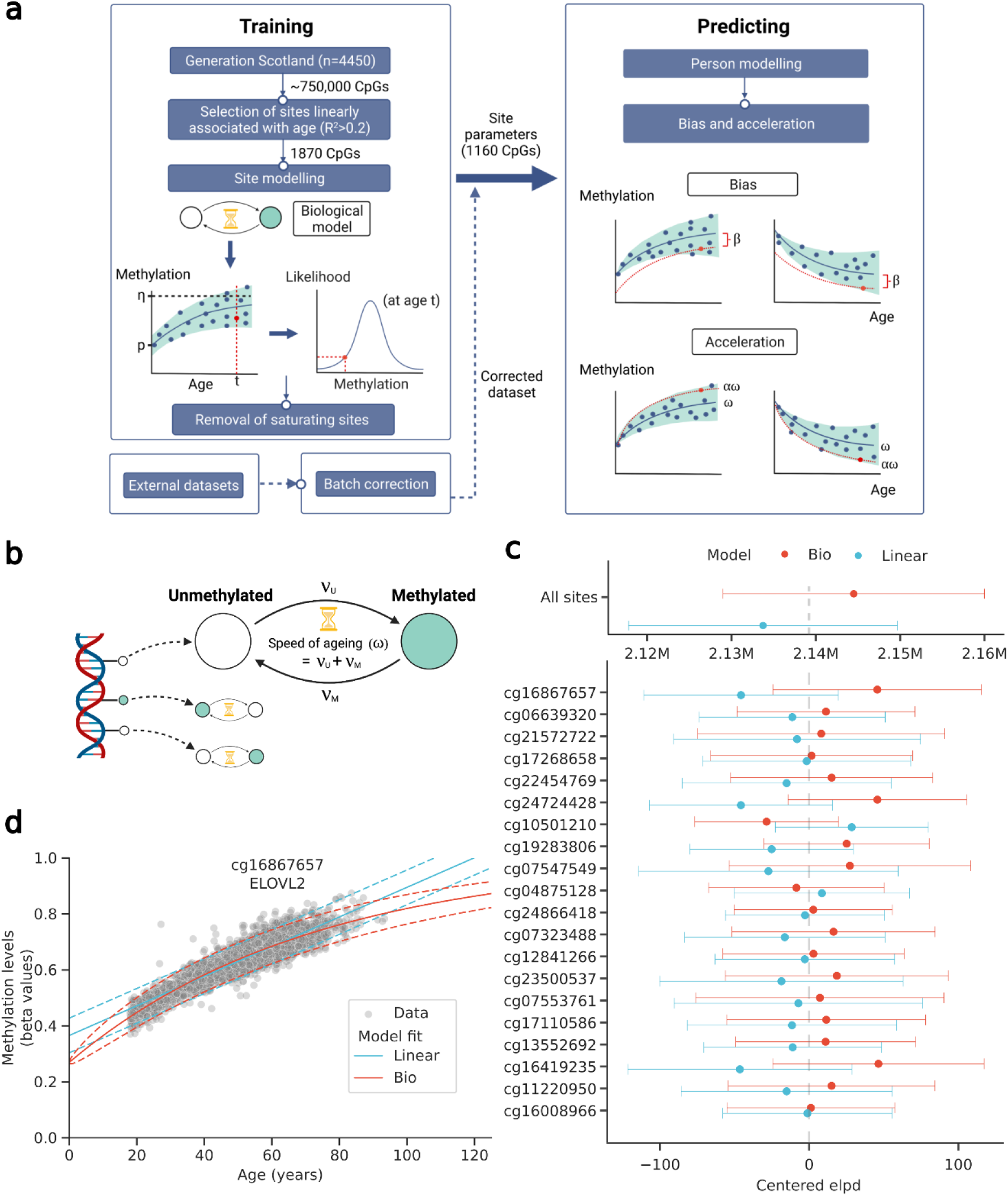
Workflow and modelling methylation on CpG sites. a. Overview of the study workflow showing how site-specific parameters are derived across the cohort and then used to calculate bias and acceleration values for each individual. There is a separate pipeline for the external datasets which does not involve training. Created with BioRender.com. b. Schematic showing the biological model’s underlying stochastic mechanics. For a single cell, multiple CpG sites can over time either methylate or demethylate with rates *ν_U_* and *ν_M_* respectively. Created with BioRender.com. c. Comparison between the linear and biological model in CpG sites. Sites are compared using the expected log-predictive density (ELPD), approximated using Pareto-smoothed importance sampling. This measure penalises the model for a higher number of parameters and gives a higher value for a model that better explains the data. For each comparison we show the mean and 2 standard deviations of the ELPD. Top plot shows the model comparison across all 1870 used for model training. The bottom plot shows the model comparison for the top 20 sites correlating with age. In the bottom plot, ELPD of each comparison has been centred at the value of zero to facilitate the display of many sites. d. Predicted dynamics of methylation levels in CpG 16867657 by the biological and linear model. Grey dots show the methylation and age of individuals in the GS_set1_ cohort. Red lines show the biological model’s mean and 95% confidence interval predictions. Blue lines show the linear model’s mean and 95% confidence interval predictions.

**Figure 3.**
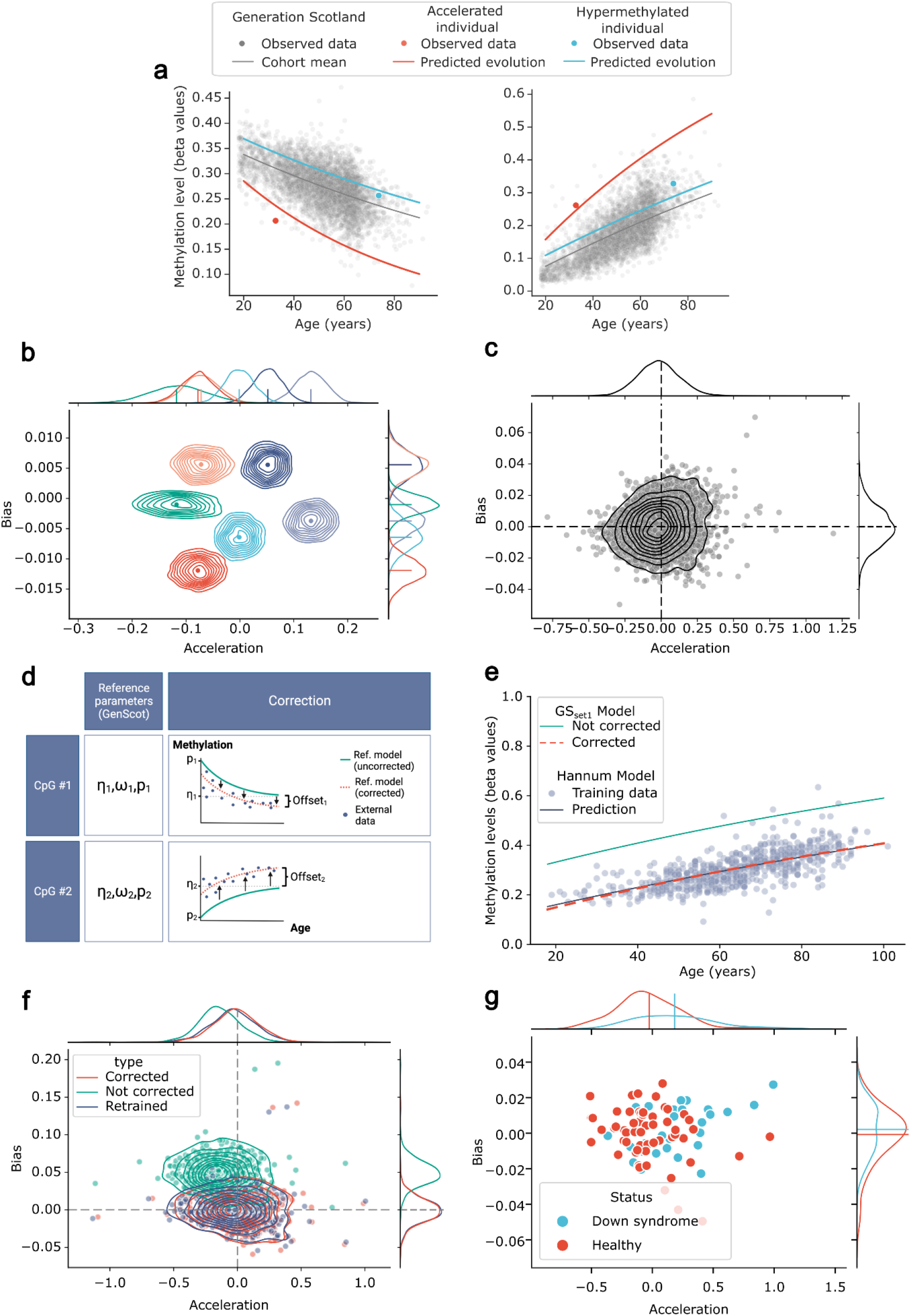
Acceleration and Bias. a. Differences between acceleration and bias in model predictions. Predicted evolution of both an accelerated but not biassed (red) individual, and of a biased but not accelerated individual (blue) in GS_set1_. The trajectories for the same individuals are shown for both sites with decreasing and increasing methylation levels with time in separate panels. Acceleration increases the *slope* of, whereas bias globally offsets, the predicted evolution of methylation levels. In grey we show the methylation and age values for the rest of individuals in GS_set1_ and the predicted mean evolution of methylation levels. b. Confidence around the inference of acceleration and bias. Joint posterior distribution of the acceleration and bias for 6 individuals in GS_set1_. Each individual is identified by a unique colour. The Maximum A Posteriori (MAP) value is also highlighted. The marginal posterior distribution of the acceleration and bias are shown in marginal plots for each individual. c. Inference results of acceleration and bias for all individuals in the GS_set1_ cohort. Marginal plots show the Kernel density estimate (KDE) of the distribution of inferred MAP values of acceleration and bias for all individuals in GS_set1_. d. Schematic illustrating how batch correction is applied on each site in an external dataset. An offset is inferred for each CpG site as the uniform shift in the predicted dynamics of our biological model that maximises the probability of observing all individuals in the external dataset. Created with BioRender.com. e. Schematic table of models used to validate the batch correction algorithm. The corrected external model denotes the model trained on GS_set1_ applied on an external dataset after batch-correction (reference parameters and offset). The uncorrected model refers to applying the trained model on GS_set1_ without batch correction on the external dataset (only reference parameters). Finally, the retrained model denotes a completely new model with parameters inferred using exclusively the external dataset. f. Visual interpretation of the effect of batch correction between GS_set1_ and the Hannum external dataset on a single CpG site. Blue dots show the methylation levels and ages of individuals in the reference GS_set1_ cohort are plotted in blue. Red dots show the methylation levels and ages of individuals in the external Hannum cohort. Blue dots show the optimal offset inferred by our batch-correction algorithm. g. Effect of batch correction on the inference of acceleration and bias values. Dots show the acceleration and bias inferred by the retrained (green), not corrected (red), and corrected (blue) models. Marginal plots show the KDE of each parameter’s distributions. h. Distribution of acceleration and bias inferred in a cohort of individuals with Down syndrome. Our model was batch corrected using the control group, and acceleration and bias were computed for both control and disease groups. Each dot represents an individual in the external cohort and is coloured according to their disease status. Marginal plots show the KDE distributions of acceleration and bias as well as the group mean.

**Figure 4.**
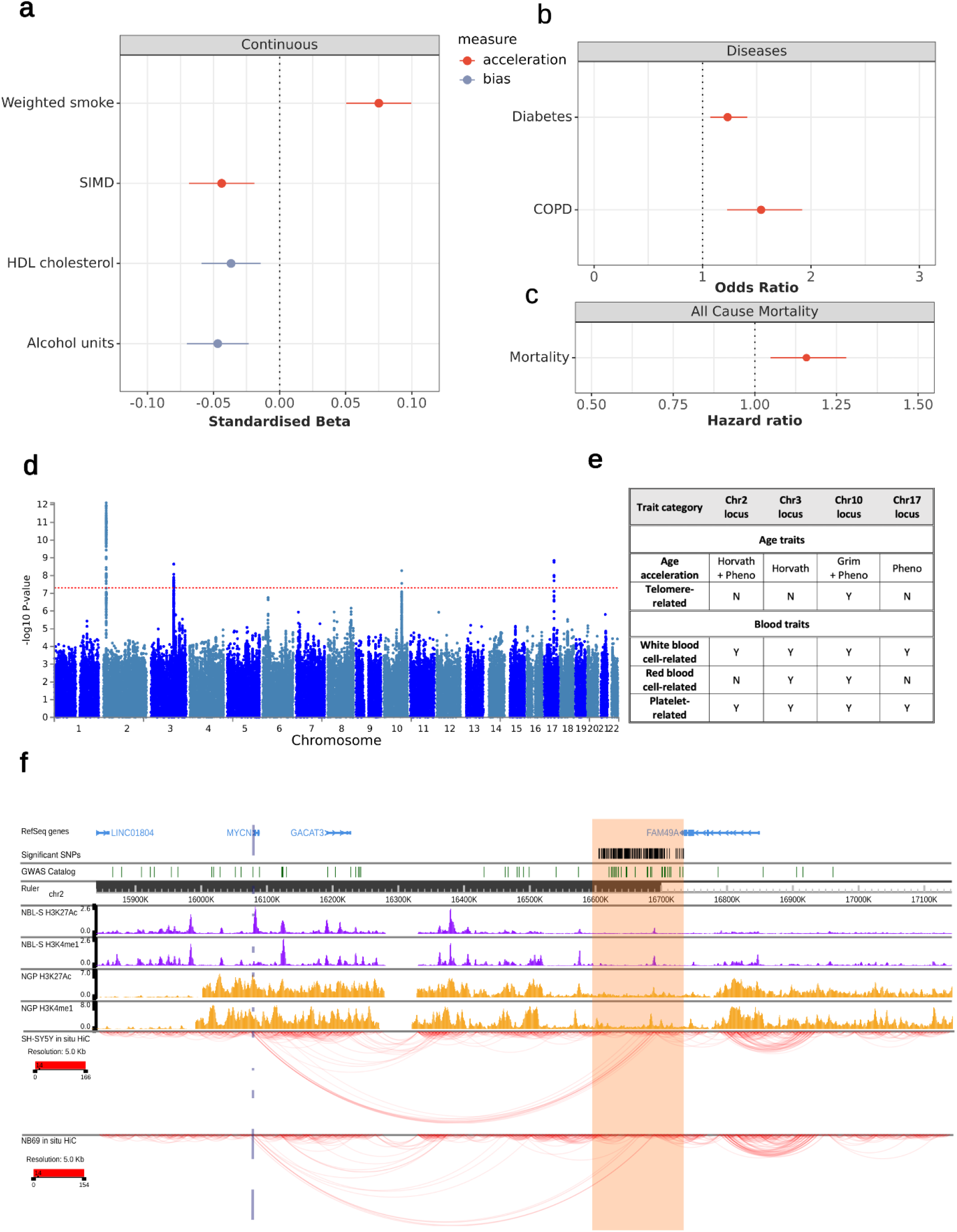
Phenotypic and genome-wide association studies. a. Associations between acceleration, bias and continuous traits. Forest plot of significant associations (FDR-adjusted P<0.05) between continuous phenotypes and age acceleration (red) and bias (purple). Results shown as standardised beta values with 95% CI from linear regressions of the form: phenotype ~ acceleration + bias + age + sex. SIMD (Scottish Index of Multiple Deprivation) b. Associations between acceleration, bias and categorical traits. Forest plot of significant associations (FDR-adjusted P<0.05) between disease phenotypes and age acceleration (red) and bias (purple). Results shown as odds ratio with 95% CI from logistic regressions of the form: disease ~ acceleration + bias + age + sex. COPD (chronic obstructive pulmonary disease). c. Associations between acceleration and all-cause mortality. Forest plot of the association between acceleration and all cause mortality (FDR-adjusted P<0.05). Result shown as hazard ratio and 95% CI from a Cox proportional hazards model of the form hazard ~ acceleration + bias + age + sex. The associations reflect an elevation of one standard deviation in the relevant measure of biological ageing. d. Manhattan plot of results from the genome-wide association analysis of age acceleration. Red dotted line indicates the genome-wide significance threshold (5×10^-8^). e. Presence of GWASCatalog trait associations of any SNPs in LD (R^2^>0.1) with the lead SNP in the four genome-wide significant genomic loci. Traits are grouped into two categories of interest – age-related and blood-related traits. f. A zoomed-out view of chromatin interactions and chromatin features in and around the genomic risk loci chr2:15841100-17142797 (highlighted in orange). Significant SNPs are shown as red vertical lines. SNPs identified by published GWAS collected in the NHGRI-EBI GWAS Catalog are shown as green vertical lines. ChIP-seq for H3K27ac and H3K4me1 in NBL-S cells and NGP cells are shown as purple histograms and orange histograms, respectively. Coloured arcs depicting In situ HiC chromatin contacts in SH-SYSY cells and NB69 cells. Demonstrating interaction of the genomic risk loci with the *MYCN* gene locus.

Taken together, this analysis showed that incorporation of lifestyle-associated naCpGs helps to improve the predictive performance of epigenetic clocks for chronological age at the expense of reducing the captured biological variability.

We next considered global biases in methylation levels. These may occur due to incomplete bisulfite conversion or as the result of an underlying biological process. The imbalanced contribution of hyper- and hypomethylating CpGs in epigenetic predictors results in over- or underestimation of the inferred acceleration (Supplementary Information Methods 2.1). To test this hypothesis we offset GS_set1_ global methylation levels both positively and negatively and applied several published epigenetic clocks to the modified data (Fig. 1g). This showed that while the degree of sensitivity to global changes in methylation varies across clocks, all displayed shifts in acceleration predictions.

Finally, acceleration is commonly defined, uniformly for all ages, as the residual between clock predictions and chronological age. However, we estimated the time-evolution of the variance of methylation levels, in all sites showing a correlation with age (R^2^>0.2), and observed an overall increase with age (Fig. 1h), as seen in other studies ^19^. This discrepancy results in an increased variance of acceleration values for older individuals (Fig 1h).

These limitations of epigenetic age predictors (summarised in Fig. 1) and the absence of a mechanistic biological foundation necessary for interpreting their outcomes motivated us to devise an alternative approach for measuring epigenetic age acceleration (Fig. 2a).

### A biological model of methylation change

Mechanism-based mathematical models offer a tangible interpretation of biological processes driving methylation patterns change over time. At a single cell level, methylation change at each genomic locus can be interpreted as a transition between unmethylated and methylated states, with rates υ_U_ and *υ_M_* respectively (Fig. 2b). This parsimonious model was first proposed in 1990 to study the clonal inheritance of CpG patterns^15^. The dynamics of methylation change over time at each site can be described with three interpretable parameters. The initial and terminal proportions of methylated cells, *p* and η, determine the directionality of the mechanism of ageing at each locus. In contrast, the total rate of transitions between cell states, ω, equates to the biological notion of speed of ageing at a single CpG site (Fig. 2a-b and Methods). This formulation allows derivation of a biologically informed notion of age acceleration, α. The dynamics associated with an accelerated ageing phenotype are described by a uniform increase in the rate of transitions, *υ_U_* and *υ_M_*, between methylation states across CpG sites (Fig. 2a). Further, this model allows inference of global hypo- or hypermethylating changes affecting the whole methylome of an individual, termed bias or β (Fig. 2a).

The linear approximation of this biological model provides a familiar interpretation of epigenetic acceleration and bias. An increase α in the speed of ageing at a single CpG site corresponds to a proportional increase of the slope in methylation trajectories. Similarly, bias corresponds to a uniform shift of the intercepts (Fig. 2a and Supplementary Information Methods).

We used the GS_set1_ cohort (n=4,450, age range 18 to 94 years) to fit our biological model and infer the parameters for each CpG site with a high age correlation (R^2^>0.2, n=1870).

To compare the fit of our approach with the linear model, we used Bayesian techniques to approximate the predictive power, expected log-predictive density, of both models in CpG sites (see Methods). Our proposed biological model outperforms linear modelling in 97% of all considered CpG sites (Fig. 2c-d) and globally across all CpG sites considered (Fig. 2c). Since the biological model predicts nonlinear dynamics, it is capable of capturing complex changes in both mean and variance that linear models fail to explain (Fig. 2d).

The model parameters inferred from the GS_set1_ cohort enabled us to quantify the extent to which each person’s methylation pattern deviates from the cohort average. This difference was explained and quantified in terms of acceleration and bias, inferred across sites for each individual separately (Fig. 3a-b and Methods). Critical for the success of this model is the capacity to disentangle the effects of both cellular mechanisms. The effects of acceleration and bias, however, can be separated because global changes in methylation vary in the same direction, acceleration is linked to the direction of methylation changes at each genomic locus (Fig. 3a). Bayesian analysis of the posterior distribution of the inferred acceleration and bias for all individuals verifies that these parameters are not correlated (Fig. 3b). We then showed that similar acceleration and bias estimates can be obtained when we consider only the top 250 CpGs with the strongest associations with age (Extended Data Fig. 2a-b and Methods).

The distributions of acceleration and bias inferred from all individuals in the GS_set1_ cohort is shown in Fig. 3c. This highlights that individuals can exhibit increased acceleration or bias, or a combination of both. We then showed that the predictions of acceleration are constant in variance across age ranges and that they are robust against global changes in methylation (Extended Data Fig 2c-d).

### Batch correction

We designed a batch correction algorithm to allow comparisons across cohorts. This enables the transferability of our model to an external cohort without the need to retrain. We do this by applying a site-specific offset to all individuals in the new cohort (see Methods) (Fig 3d). We tested the validity of our batch correction algorithm on the Hannum *et al*. cohort ^2^. In Fig. 3e (and Extended Data Fig. 2e) we provide a visual comparison of the effect of our batch correction algorithm on the site showing the largest offset. We showed minimal differences in predictions between our batch-corrected model and a fully retrained model on the external dataset (Fig 3f and Extended Data Fig 2f).

To further validate our batch correction algorithm, we investigated its applicability to datasets with limited sample sizes. We used publicly available data of a Down syndrome cohort (n=58 control, n=29 disease)^20^. Despite the dataset’s small number of participants, Down syndrome was associated with an increase of 0.7 standard deviations in epigenetic age acceleration (p=0.002, see Methods).

### Acceleration is associated with lifestyle factors and health outcomes

Next, we tested for association of age acceleration or bias with disease phenotypes and genetic variants. The following analyses were conducted on the combined subsets of Generation Scotland (n=7028) - GS_set1_ and the batch corrected subset GS_set2_, (n=2578, unrelated to each other and to GS_set1_, and processed in separate experimental batches).

We performed linear regression analyses to quantify associations between acceleration and disease phenotypes while controlling for age and sex (Fig. 4a-c, Supplementary Table 1). All of the significant results have passed the false discovery rate (FDR) adjusted value threshold. This revealed significant (FDR-adjusted *P*<0.05, see Methods) associations between age acceleration and four phenotypes. Increased acceleration was associated with higher levels of tobacco smoking and with greater deprivation (as measured by the Scottish Index of Multiple Deprivation) (Fig. 4a). Increased acceleration was also associated with a number of self-reported prevalent disease outcomes, namely diabetes and chronic obstructive pulmonary diseases (Fig. 4b). Further, a Cox proportional hazards regression analysis showed an association between acceleration and all-cause mortality (Fig. 4c, hazard ratio = 1.158 per standard deviation increase of acceleration; 95% confidence interval 1.048, 1.280; FDR-adjusted *P*=1.34×10^-2^). Similarly, increased bias was shown to be associated with two phenotypes, namely lower alcohol consumption and HDL cholesterol levels. These associations were also recapitulated to a large extent when computed separately on both cohorts GS_set1_ and GS_set2_ (Extended Data Figure 3a-c, Supplementary Table 1).

These results highlight that our measure of increased speed of ageing captures biologically meaningful age-related traits. Additionally, the lack of overlap between associations found with acceleration and bias suggests that these two parameters capture distinct biological phenomena (Fig. 4a).

### GWAS of acceleration implicates age-related genomic loci

We then conducted a genome-wide association analysis to identify genetic variants that are associated with acceleration. We found 218 SNPs at genome-wide significance (p<5×10^-8^) clustered in four genomic regions on chromosomes 2, 3, 10 and 17 (Fig. 4d). Acceleration-associated SNPs clustered into two main categories of interest: age- and blood-related traits (Fig 4e). All four genomic regions contain SNPs associated with age acceleration as measured by other ageing clocks^21^ (Horvath, PhenoAge, or GrimAge), and SNPs within the chromosome 10 region (chr10: 101271789 - 101589328) are genome-wide associated with telomere length. All regions also contain SNPs associated with numerous blood-related traits.

We then performed downstream functional analysis to link SNPs with genes by overlapping the genomic regions with PBMC promoter capture Hi-C data^22^ and PBMC ChIP-seq data from the ENCODE project^23^ (Extended Data Fig 3d). The strongest association was found for a locus in chromosome 2 with a distal interaction with the *MYCN* promoter region. *MYC*, a paralogue *MYCN*, has been convincingly shown to play a crucial role in determining lifespan and shaping various aspects of health and well-being in mammals^24^. We validated this long-range interaction in several datasets, including *in situ* HiC data from primary cancer cell lines of *MYCN*-driven neuroblastoma, in which we can see a pronounced interaction (Fig. 4f). Furthermore, the found promoter-interacting region overlaps with H3K27ac and H3K4me1 histone marks, suggestive of enhancer activity. A SMC1 Chromatin Interaction Analysis with Paired-End Tag (ChIA-PET) experiment showed that SMC1 sites located at the enhancer/snp and *MYCN* promoter are interacting^25^ (Extended Data Fig 3e). Once again, this strengthens the case for a physical promoter-enhancer intersection. This link was reinforced by the association of this locus with various cleft lip/palate traits (Extended Data Fig. 3e), for which MYCN deficiency is a known risk factor^26^. ^27^

## Discussion

The emergence of epigenetic age predictors was a watershed moment in geroscience, propelling the measurement and concept of biological ageing into a quantitative era.

Our study attempts to integrate biological mechanisms and quantitative principles to measure biological age. Firstly, we identify and address technical limitations in current epigenetic predictors trained on chronological age. Secondly, we provide a biologically tangible interpretation of our outputs. This has been a long-held criticism inherent to previous clocks that has hindered the progress in accurately defining the concept of epigenetic speed of ageing and in studying the functional mechanisms underlying this process. This study provides a biological interpretation of the speed of ageing, at a single-CpG level, defined as the total rate of chemical reactions occurring in cells that result in changes between their methylation states. Two notions central to methylation dynamics naturally emerge in this model: age acceleration, defined as the proportional increase in the speed of ageing across sites, and bias, global changes in methylation. Our study showed that both acceleration and bias are associated with distinct disease phenotypes. Additionally, we used a novel batch-correction algorithm, which can be used to improve comparison between cohorts or to explore smaller datasets.

We consider our model to be parsimonious, in that it uses the minimal set of parameters required to accurately recapitulate the observed methylation dynamics. However, a limitation that arises from this approach is the simplification of the mechanism by which cells can gain or lose methylation. Inclusion of enzymatic processes in a more complex model could offer an alternative interpretation of ageing on an enzymatic level (also discussed in Supplementary Information Methods 1.3).

Additionally, our model does not account for cell count compositions. However, our analysis of the methylation dynamics observed in CpG sites challenges the widespread hypothesis that methylation changes with age are predominantly influenced by variations in cell-type composition. If this were the case, we should observe clustered patterns of speed of ageing across different sites corresponding to different cell types, whereas we observe a continuum. Further, the genomic locus that correlated strongest with age in Generation Scotland and other studies^6^ “cg16867657” (*ELOVL2*), shows a dynamic range in methylation that is inconsistent with cell-type composition changes at old age. Although these remarks are not conclusive, the hypothesis that changes in cell-type composition cannot account for the observed trends in methylation changes at clock CpG sites has already been convincingly argued in previous studies ^28^.

The biological relevance of acceleration measured by our model was substantiated using both phenotype association analysis (in combined and independent subsets of GS) and GWAS. Acceleration demonstrated significant correlations with various phenotypes and disease burdens, including diabetes and COPD. These are leading causes of death in high-income countries^29^, and is reflected in our analysis by showing increased risk of mortality associated with increased acceleration. Interestingly, we see the strongest overlap of our GWAS associations not with the outcomes of first generation clocks, but with those reported by the composite clock PhenoAge. Although replication in other cohorts should be sought, this highlights that our proposed model captures biologically relevant information.

Downstream functional analysis of the SNPs associated with age acceleration found genetic variants inside a distal enhancer region for the *MYCN* gene. *MYC*, the paralogue to *MYCN*, has been associated with ageing in a comprehensive study suggesting that the activity of *MYC* plays a crucial role in determining lifespan and affecting various aspects of health and well-being in mammals^24^. *MYCN* as a general cell proliferator in development and the contrasting role in age acceleration make it a candidate gene for antagonistically pleiotropic effects^30^.

## Methods

### Generation Scotland

#### Overview

Generation Scotland (GS) is a family-based cohort consisting of individuals, aged 18 to 99 years, living across Scotland. Recruitment took place between 2006 and 2011. The cohort encompasses 5,573 families with a median family size of 3 (interquartile range=2–5 members; excluding 1,400 singletons without any relatives in the study). Full details on the cohort and baseline data collection have been described previously ^31,32^.

#### DNA methylation

Genome-wide DNA methylation was measured from blood samples using the Illumina Infinium HumanMethylationEPIC BeadChip at >850,000 CpG sites. The methylation profiling was carried out in two sets, here referred to as Set 1 and Set 2. Set 1 consisted of 4,450 unrelated individuals, who were also unrelated to individuals in Set 2. Set 2 consisted of 2,578 individuals who were genetically unrelated to each other at a relatedness threshold of < 0.025. Poor performing probes, X/Y chromosome probes and participants with unreliable self-report data or potential XXY genotype were excluded. Full details of DNA methylation quality control steps are detailed under Supplementary Information Methods 6.1.

#### Genotyping

Generation Scotland samples were genotyped using the Illumina Human OmniExpressExome-8v1.0 and 8v1.2 BeadChips, and processed using the Illumina Genome Studio software v2011 (Illumina, San Diego, CA, USA). Quality control steps are outlined in full under Supplementary Information. Duplicate samples, samples with genotype call rate < 0.98 or outlier values on PCA of genotype data were excluded. SNPs with a call rate < 0.98, minor allele frequency ≤ 0.01 and Hardy-Weinberg equilibrium test with p ≤ 1 x 10-6 were also removed. Genotype data were imputed using the Haplotype Research Consortium (HRC) dataset and ~24 million variants were available for analyses ^33,34^. There were 7023 individuals with genotype and methylation data. Full details of genotyping quality control steps are detailed under Supplementary Information Methods 6.2.

#### Phenotyping and health record linkage

GS participants self-reported health and lifestyle data at the study baseline, including lifetime and family history of approximately 20 disease states. Over 98% of GS participants consented to allow access to electronic health records via primary and secondary care records (i.e. Readv2 and ICD codes). Data are available prospectively from the time of blood draw, yielding up-to-15 years of linkage. Information on mortality and cause of death are updated via linkage to the National Health Service Central Register, which is provided by the National Records of Scotland (data correct as of March 2022).

#### Ethics

All components of GS received ethical approval from the NHS Tayside Committee on Medical Research Ethics (REC Reference Number: 05/S1401/89). GS has also been granted Research Tissue Bank status by the East of Scotland Research Ethics Service (REC Reference Number: 20-ES-0021), providing generic ethical approval for a wide range of uses within medical research.

### Associations of other epigenetic clock CpGs with age and smoking

CpG sites were considered naCpGs linear regressions of the form *meth* ~ *age* showed a coefficient of determination R^2^<0.1. To compute associations between methylation levels and smoking, we created the “weighted smoking” variable which combined information on both how much a participant smoked at any point of in its life, *norm_smoke* = (1 + *packs/year*), and the current smoking status, *curr_smoke*, categorically defined by: 1-currently smoking, 2-stopped within 12 months, 3-stopped more than 12 months ago, 4-never smoked. More precisely we assumed that the effect of smoking is proportional to *norm_smoke* and fades exponentially with their smoking status, that is *weighted_smoke = norm_smoke/exp(curr_smoke*). In binary definitions of smoking, we used a *weighted_smoke* cutoff of 0.25 based on the observation from the cohort-wide histogram that the vast majority of people clustered below 0.25, with a long tail of heavier smokers above 0.25. When calculating smoking associations for the figures, we used this binary definition of smoking and reported the coefficient of smoking from linear regressions for each CpG of the form: *meth ~ weighted-smoke* + *age* + *sex*.

### Bootstrapping linear regression models

To show the decline in associations with increasing training size cohorts, we selected random training cohorts with set sizes and test sets (n=890, or 20% of GS_set1_) from GS_set1_. We trained Lasso models with 5-fold cross validation to allow optimisation of the selected CpG sites and regularisation levels.

We then computed acceleration as the residual from the model prediction and chronological age and inferred its association with *weight_smoke* on the test set.

We bootstrapped linear regression models with increasing proportions of smokers (categorically defined as *weight_smoke >* 0. 25) similarly. Here training sets had a fixed size (n=700) limited by the amount of smokers.

### R^2^ filter of CpG sites

To avoid numerical errors during model fitting in GS_set1_ we discarded CpG sites with NaN methylation values (n=6) and replaced methylation levels of 0 and 1 by 0.0001 and 0.9999, respectively. Next, we fitted linear regression.

We then fitted linear regression models of the form *meth ~ age* on the remaining sites (n=773854).To maximise the age correlation of sites with ageing used for model training, and to avoid the presence of naCpG sites in our model, we filtered all CpG sites showing a low coefficient of determination, R^2^ <0.2. The remaining sites (n=1870) were taken forward for model comparison.

### Biological modelling of methylation dynamics at a single CpG site level

We now present an overview of the key results necessary to understand the derivation and interpretation of the proposed mathematical model of methylation dynamics. An exhaustive description and precise mathematical derivations can be found in the Supplementary Information.

To model the evolution of methylation dynamics with time, in a single CpG site, we considered a minimal model of transitions between two states. The proposed model directly relates to the early work of Markov in 1905 and was used for the first time to model the dynamics of methylation in ^15^, and more recently in the context of ageing in ^16^. In this model cells can be in either an unmethylated or methylated state, *U* and *M* respectively, and can transition from one state to the other at rates *ν_U_* and *ν_M_* (Fig. 2B).

The mean evolution of this system as a function of time, *t*, is given by

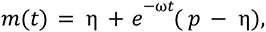

where ω = *ν_U_* + *ν_M_* corresponds to the total rate of transitions, *p* the proportion of methylation cells at initial time *t =* 0, and η *= ν_U_/ω* to the proportion transitions associated with a gain in methylation (Fig. 1A). Notice that the definition of speed of ageing at a single CpG site, ω, emerges naturally, while the directionality of these changes is independent from the speed, given by the initial state of the system *p* and the final state η.

The evolution of the variance of this system can be similarly described as a function of time,

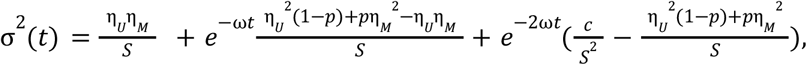

where S corresponds to the system size and *c* the initial variance at the cell level.

Although the distribution for the evolution in time of this system is not analytically solvable, it is well approximated by a beta function conditional on the mean and variance derived above. We can therefore model the probability of seeing a methylation level *m_i_* in site *i* in an individual aged *t* by

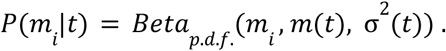

We use this probability definition to find the optimal parameters for each site using all cohort methylation observations. That is the parameters that maximise

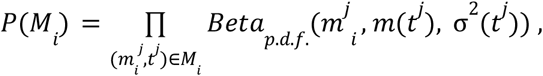

where 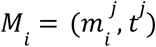 corresponds to the pair of methylation value and age of each individual in the cohort. We approximated the full posterior distribution, using a Markov chain Monte Carlo algorithm, and extracted the maximum a posteriori values of the parameters for each site using the PyMC python package.

To ease interpretation of our results, we take the log2 transformation of the original acceleration parameter, centering the cohort average at 0, and report the maximum a posteriori (MAP) estimates for both acceleration and bias.

### Model comparison

Similarly, we fitted a probabilistic linear model of the form

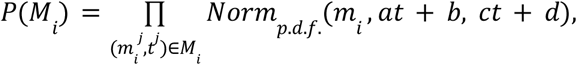

and inferred the posterior distribution and performed a model comparison using the PYMC python package. Model comparison approximated the expected log-predictive density (ELPD) of both models on each site using an approximation of leave-one-out cross validation (LOO-CV) based on the Pareto-smoothed importance sampling (PSIS). The higher ELPD value on each site highlights the favoured model to explain the observed dynamics. Since the evolution of methylation across CpG sites is assumed to be independent of each other, we compute the overall ELPD for either model by summing the reported value on all fitted sites.

### Saturation filtering

Further quality control measures were taken to ensure that only sites that capture the full expected dynamics of methylation changes with time are retained for further modelling (Extended Data Fig. 2a).

We observed that the methylation dynamics are constrained between 0 and 1. This reduces the possibility to observe deviations from the mean in sites approaching these boundaries. We therefore dropped sites for which the 95% confidence interval predicted by our model reached a threshold of either 0.005 and 0.95, at either birth or 90 years. 204 sites were observed to display this phenomenon.

Further, we noticed that the above mentioned phenomenon might occur at different thresholds, due to different batch correction processes. However, in these sites, we should see a plateau in the evolution of the methylation dynamics. We therefore computed the derivative of the mean methylation dynamics for each site, m′(t) = *ωe* ^−ωt^(η – p), and valued it at *t =* 90. This value is a measure of our distance to steady state at old age, *ds = m′(90*), at old age. We then filtered all sites with low distances to steady state, *ds* < 0. 001 (n=654).

Overall a total of 1160 sites were taken forward for acceleration and bias modelling.

### Acceleration and bias model

First, we define the probability of observing a methylation pattern in an individual by comparing its methylation levels to those obtained from cohort fitting on each CpG site. That is, if an individual of age *t* has methylation levels {m_i_}_i∈I_ on a set of CpG sites *I*, we define its probability of observation as

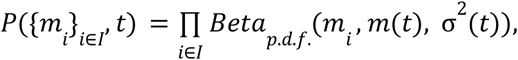

where *m(t*) and σ^2^ (*t*) are defined as above, using the MAP values for all site parameters in the model obtained from the cohort fitting. We then consider two extra parameters α and β in our model. Mathematically, we multiply the speed of reactions ν_U_ and *ν_M_* uniformly across all sites by a factor α. Notice that this translates in a proportional increase of the total speed of reaction, that becomes *αω*, for each CpG site *i*. Parameter α therefore represents a uniform increase in the average speed of ageing across all measured sites. Analogously, parameter *β* modifies uniformly all sites shifting the mean intercept, but not disturbing the expected speed of change. These two parameters modify the mean evolution of methylation on for each site as follows:

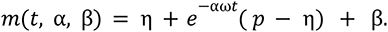

Details on how these parameters modify the evolution of the variance can be found in the Supplementary Information Methods Section 4. We can then compute the probability of observing an individual conditional on a given acceleration and bias

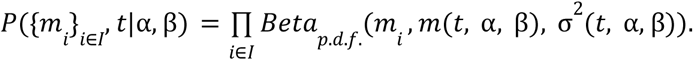

We then infer the posterior distribution of parameters α and β and compute the maximum a posteriori values that maximise the probability of observing the methylation levels in an individual across sites using the PYMC python package.

### Downsampling

To determine the optimal amount of CpG sites included for the inference of the acceleration and bias of each individual we conducted a downsampling experiment. We inferred acceleration and bias using increasing numbers of CpG sites, ordered decreasingly according to their absolute correlation with age. We then computed the absolute difference between the inferred values for each fit to infer the stability of our predictions as a function of the sites included in the model. We found that there are no benefits in including more than 250 sites (Extended Data Fig. 1b).

### Global hypo-hyper methylation test

We tested the robustness of our accelerated person model by transforming the training GS_set1_ cohort, by applying a global hypo or hyper methylation to all the data points. Then we inferred the acceleration and bias values on this transformed dataset using the site parameters inferred from the not transformed dataset (Extended Data Fig. 1d). As a benchmark, we also predicted the age acceleration using the Horvath epigenetic clock which shows little robustness against these type of methylation transformations.

### Site batch correction

When applying our model to external datasets, we do not retrain the biological model on the new data. Instead, we applied an offset to the evolution of the mean *m(t*), for each site, inferred using our reference dataset GS_set1_. Mathematically, for each site we find the offset, o_i_, that maximises the probability of observing

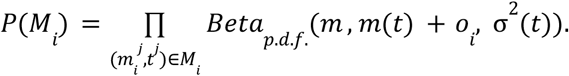

The full mathematical description of our algorithm can be found in Supplementary Information Methods section 5. We benchmarked our results on the Hannum dataset by comparing a fully retrained model on this dataset and the model trained on GS_set1_ applied both with and without batch correction. We found that offset does not have a clear directionality across sites and should therefore be applied separately for each CpG site considered in the model.

### Down Syndrome analysis

We used the control group in the Down syndrome dataset ^20^ to fit our batch correction algorithm and inferred acceleration and bias parameters for the control (n=58) and disease (n=29) groups. We then fitted a logistic regression model of the form *disease* ~ *scale(acceleratiori*).

### Association Analysis

We utilised GS_set2_ and GS_set1_’s linked health data records (n=7028) to perform association studies grouped into three categories associated with the nature of the studied traits: continuous, categorical and survival.

For each continuous phenotype, outlier values that were more than 3.5 standard deviations away from the mean were removed prior to analyses. We also applied a log transformation to the body mass index to normalise the data. Finally, we used log transformations on the alcohol consumption to reduce skewness in their distributions. This was done by applying a log(units+1) transformation. We quantified the continuous trait of smoking using our developed *weighted_smoke* parameter (described above).

### Statistical analysis

Linear regression models of the form *pheriotype ~ scaled(α*) + *scaled(β*) + *age* + *sex* were used to examine the association between continuous traits and the acceleration and bias measurement. Scaling is performed to standardise the data to a mean of zero and a variance of one. Logistic regression was used to test the association between categorical disease phenotypes and acceleration and bias. We fitted a Cox proportional hazards regression to examine whether our measures of biological ageing were associated with incidences of all-cause mortality. All of the results were corrected using the FDR method.

### GWAS and functional analysis

Genotype_phenotype association analyses were performed using a linear mixed model GWAS implemented in fastGWA GCTA ^35^. 7023 overlapping individuals with non-missing genotype and phenotype data were included. Variants with MAF < 0.01 or missingness rate > 0.1 were excluded from the analysis. Preliminary functional annotation and was performed using FUMA ^36^. Genomic risk loci were defined around significant variants (p<5 × 10-8) and included all variants correlated (R2 > 0.6) with the lead variant. We used LDtrait ^37^ to search for phenotypes associated with SNPs in linkage disequilibrium with the four lead SNPs (R2>0.1) and within 1Mb. LDtrait reports association data from the GWASCatalog ^38^. All coordinates in this study were based on human reference genome assembly GRCh37/hg19 (see URL in Framework Implementation). Gene annotations were based on gencode annotation release 39 (see URL in Framework Implementation).

## Framework Implementation

The inference of the posterior distribution of model parameters was implemented in Python version 3.9 with dependencies on PyMC 5.0.2 ^39^, Numpy 1.24.1 ^40^, Anndata 0.8.0 ^41^ and Pandas version 1.5.3 ^42^. Cox proportional hazards regression was done using the survival package version 3.4.0 ^43^, while linear and logistic regression were done using the stats base package under R base version 4.2.2. GWAS summary statistics was generated using GCTA version 1.93.2. Functional analysis of the GWAS results was done using FUMA version 1.5.1.

Human reference genome assembly used for GWAS: http://www.ncbi.nlm.nih.gov/assembly/2758/. Gencode annotation researle 39: https://www.gencodegenes.org/human/release_39.html.

## Data and code availability

All code used in this manuscript is available at https://github.com/zuberek/NSEA

According to the terms of consent for Generation Scotland participants, access to data must be reviewed by the Generation Scotland Access Committee. Applications should be made to

## Acknowledgements

The authors thank all of the participants of the Generation Scotland: Scottish Family Health Study as well as study team members for their previous and ongoing contribution to this study.

GS received core support from the Chief Scientist Office of the Scottish Government Health Directorates (CZD/16/6) and the Scottish Funding Council (HR03006). DNA methylation profiling of the GS samples was carried out by the Genetics Core Laboratory at the Wellcome Trust Clinical Research Facility, Edinburgh, Scotland, and was funded by the Medical Research Council UK, the Brain & Behavior Research Foundation (Ref: 27404) and the Wellcome Trust (Wellcome Trust Strategic Award ‘STratifying Resilience and Depression Longitudinally’ ((STRADL) Reference 104036/Z/14/Z)). We thank C. P. Ponting for critical reading of the manuscript. We thank Nelly Olova and all members of the Chandra lab for their input.

J. D. is supported by the United Kingdom Research and Innovation (grant EP/S02431X/1), UKRI Centre for Doctoral Training in Biomedical AI at the University of Edinburgh, School of Informatics. E.J.Y. is funded through core funding to the MRC Human Genetics Unit (MC_ST_U16003)

E.L.C. is a cross-disciplinary postdoctoral fellow supported by funding from the University of Edinburgh and the Medical Research Council (MC_UU_00009/2). T.C. is supported through a Chancellor’s Fellow at the University of Edinburgh and the MRC Human Genetics Unit.

S.J.C.C is supported by the Wellcome Trust Hosts, Pathogens & Global Health Programme (grant number: grant.226831/Z/22/Z).

## Author information

These authors contributed equally: Jan Dąbrowski and Emma J. Yang

These authors contributed equally: Eric Latorre-Crespo and Tamir Chandra

### Contributions

E.L.C. and T.C conceived and supervised the study. J.K.D., E.J.Y, S.J.C.C, E.L.C and T.C. wrote the manuscript. J.K.D. and E.L.C. developed the modelling framework. J.K.D., E.J.Y., S.J.C.C., D.J.S. and T.C. conducted data analysis. R.F.H., D.L.M and R.E.M. curated Generation Scotland, gave access to data and advised on the study.

## Ethics declarations

R.E.M. is an advisor to the Epigenetic Clock Development Foundation. R.E.M. and R.F.H. are advisors to Optima Partners.

## Additional information

Supplementary Information is available for this paper.

## Extended Data

**Extended Data Figure 1.**
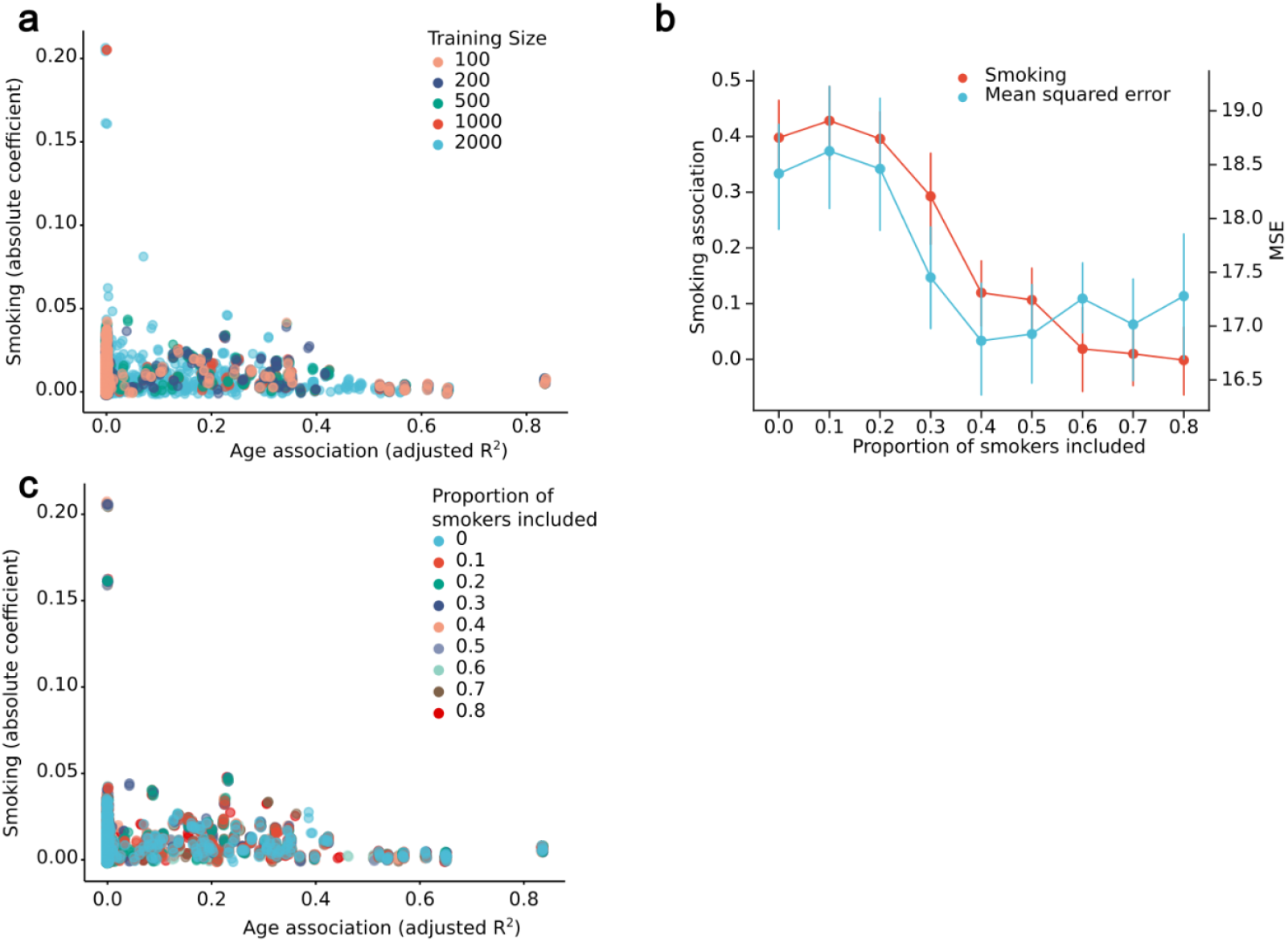
a. Comparison between the association of methylation levels with smoking and age on each CpG site included in a training size bootstrap experiment. Each point corresponds to a site included in a clock, coloured by the size of the dataset used to train the epigenetic predictor and sized proportionally to the smoking coefficient value. Age association displayed as adjusted R^2^ from linear regressions for each CpG of the form: methylation ~ age. Smoking association shown as the absolute value of the coefficient of smoking (dichotomised as a weighted smoking value of greater than 0.25) from regressions for each CpG of the form: methylation ~ smoking + age + sex. Weighted smoking is defined as log(1 + pack years)/exp(ever smoke). Points are displayed with a random jitter to avoid overlap. b. Acceleration obtained from bootstrapped lasso linear regressions trained on chronological age trained on datasets of increasing proportions of smokers and a fixed size (n=700). Training cohorts were randomly sampled from the GS_set1_ dataset. The results were computed on random test sets (n=2000). The red line, associated to the left y-axis, shows the average, and 95% confidence interval, association between the predicted age accelerations with smoking, given training cohort size. The blue line, associated to the right y-axis, shows the average, and 95% confidence interval, mean squared error in the prediction of chronological age. c. Comparison between the association of methylation levels with smoking and age on each CpG site included in a bootstrap experiment of the effect of the proportion of smokers. Each point corresponds to a site included in a clock, coloured by the size of the dataset used to train the epigenetic predictor and sized proportionally to the smoking coefficient value. Age association displayed as adjusted R^2^ from linear regressions for each CpG of the form: methylation ~ age. Smoking association shown as the absolute value of the coefficient of smoking (dichotomised as a weighted smoking value of greater than 0.25) from regressions for each CpG of the form: methylation ~ smoking + age + sex. Weighted smoking is defined as log(1 + pack years)/exp(ever smoke). Points are displayed with a random jitter to avoid overlap.

**Extended Data Figure 2.**
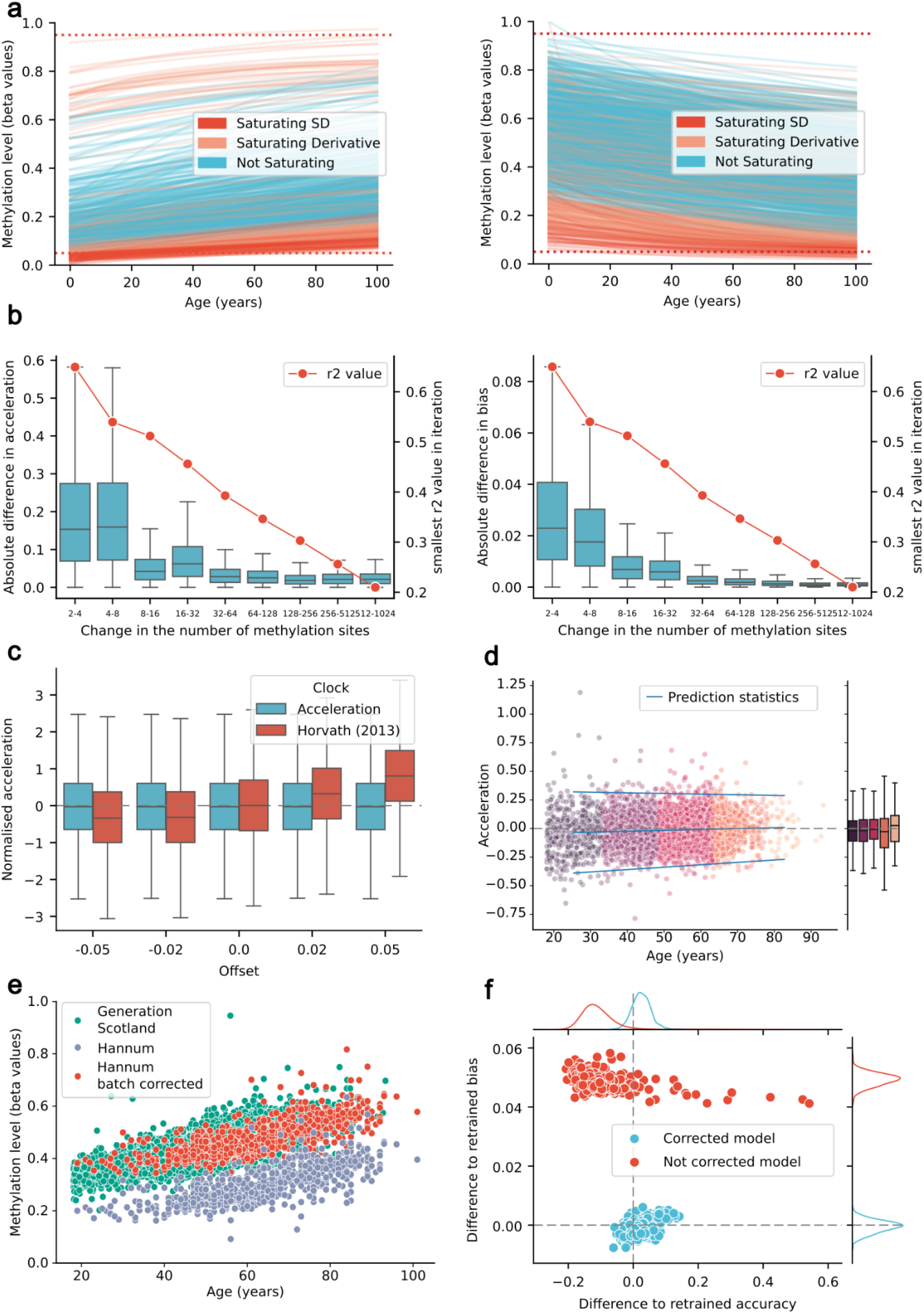
a. Filter of saturating sites showing increasing (left panel) and decreasing (right panel) methylation trends. Each line shows the predicted time-evolution of the mean methylation level of a single CpG site. Red lines represent sites saturating based on the mean value at the age of 90, orange lines represent sites saturating based on the derivative value of our model prediction at the age of 90. Blue lines indicate sites that have not been filtered and are taken forward for acceleration and bias fitting. Horizontal dashed lines at values 0.05 and 0.95 are included, to show the threshold for the mean saturation filter. b. Effect of increasing the number of CpG sites on the inferred acceleration (left) and bias (right) for all participants in the GS_set1_ cohort. Acceleration and bias were fitted using exponentially increasing numbers of CpG sites decreasingly ordered in terms of the coefficient of determination, R^2^, between the methylation levels and age of participants in each site. Boxplots show mean and exclusive interquartile range of the absolute difference in consecutive inferences of acceleration and bias. Additionally, a the lowest R^2^ value included in each model is shown in red for each iteration, with values shown on the second Y-axis c. Effect of batch correction on the inference of acceleration and bias values in the Hannum dataset. Dots show the difference from the acceleration and bias using the corrected (blue) and not corrected (red) models to that inferred using the retrained model. Marginal plots show the KDE of each parameter’s distributions. d. Impact of different global offsets, affecting all CpG sites uniformly, on the accelerations inferred by our model and Horvath’s epigenetic predictor. Accelerations are computed without applying the batch correction. Box plots show the median and exclusive interquartile range of the accelerations inferred for all individuals in the GSset1. Acceleration values are scaled by the standard deviation observed in GSset1 to facilitate a direct comparison. In the biological model, bias absorbs the effect of global changes in methylation and results in a stable acceleration measure. e. Age-stability of the inferred accelerations in individuals of GS_set1_. Each dot corresponds to the acceleration and age of individuals in the GSset1 cohort. In blue, we show a linear regression of the mean and 2 standard deviations of the predicted accelerations, computed using age bins reflected by the change in colour of points. The right marginal box plot shows the median and exclusive interquartile range of the predicted accelerations by bin.

**Extended Data Figure 3.**
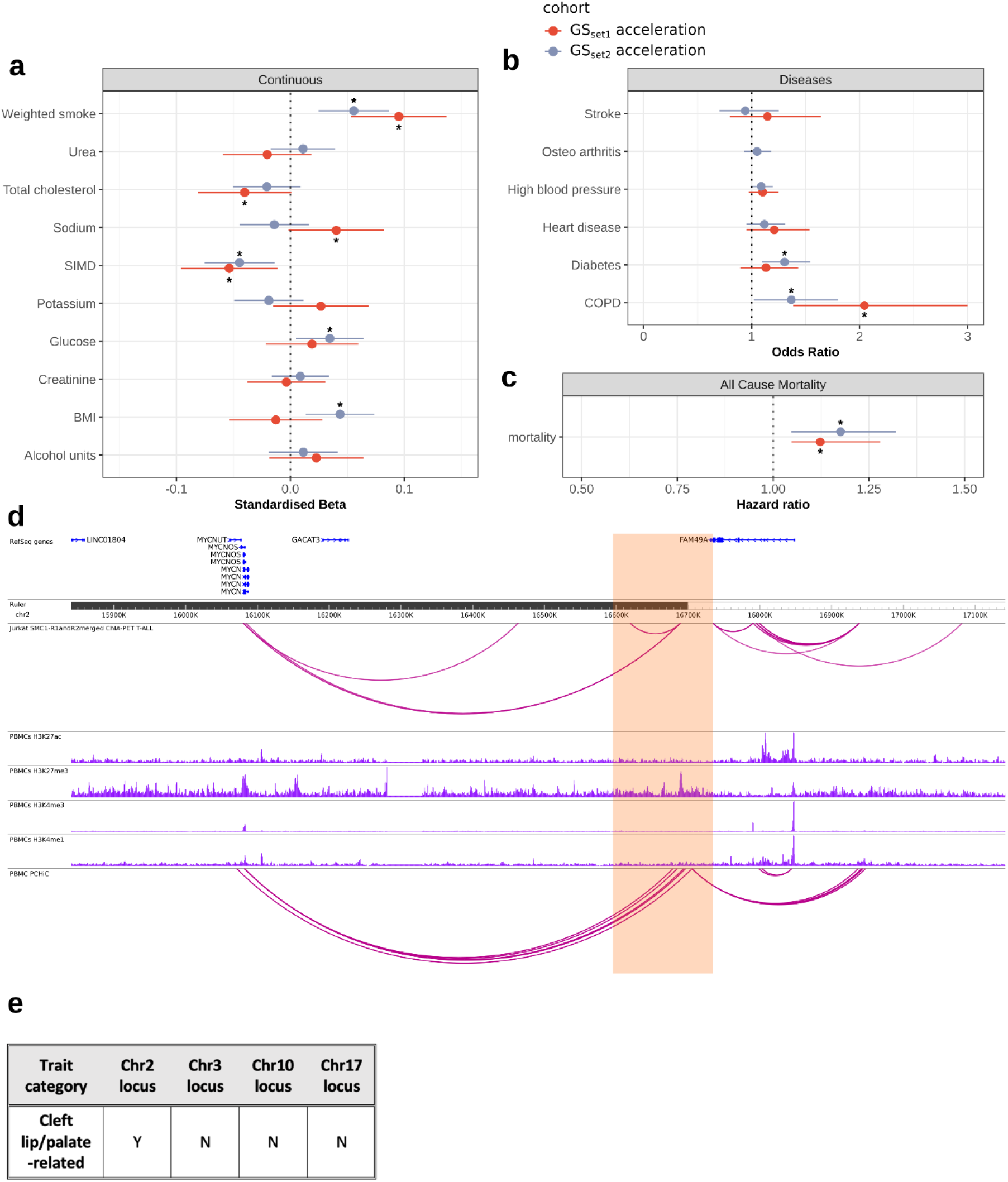
a. Associations between GS_set2_ acceleration, GS_set1_ acceleration and continuous traits. Forest plot of associations between continuous phenotypes and age acceleration from GS_set2_ (orange) and GS_set1_ (purple). Results shown as standardised beta values with 95% CI from linear regressions of the form: phenotype ~ acceleration + bias + age + sex. Significant associations (FDR-adjusted P<0.05) are highlighted with asterisks. b. Associations between GS_set2_ acceleration, GS W3 acceleration and categorical traits. Forest plot of significant associations (p<0.1 for GS_set2_ and p<0.05 for GS_set1_) between disease phenotypes and age acceleration from GS_set2_ (orange) and GS_set1_ (purple). Results shown as odds ratio with 95% CI from logistic regressions of the form: disease ~ acceleration + bias + age + sex. Significant associations (FDR-adjusted p<0.05) are highlighted with asterisks. c. Associations between acceleration and all-cause mortality. Forest plot of the association between acceleration and all-cause mortality (p<0.05). Result shown as hazard ratio and 95% CI from a Cox proportional hazards model of the form hazard ~ acceleration + bias + age + sex. The associations reflect an elevation of one standard deviation in the relevant measure of biological ageing. Significant associations (FDR-adjusted p<0.05) are highlighted with asterisks. d. A WashU genome browser showing genetic and epigenetic information within and around the genomic risk loci chr2:15841100-17142797 (highlighted in orange). Blue histograms indicate PBMC ChIP-seq data, while top purple arcs depict SMC1 ChIA-PET data in T-ALL Jurkat cells, and bottom purple arcs represent promoter capture HiC data in PBMC cells. e. Presence of GWASCatalog cleft lip/palate trait associations of any SNPs in LD (R^2^>0.1) with the lead SNP in the four genome-wide significant genomic loci.

