## Supplementary Information Methods for "Probabilistic inference of epigenetic age acceleration from cellular dynamics"

### Probabilistic models of epigenetic aging

February 2023

Here we develop a probabilistic model of epigenetic ageing, based on a biological system of methylation changes. This yields a method to compute the probability of observing the methylation profile of an individual across CpG sites within a cohort of individuals. For this, we turn our attention to modelling the biological mechanism of methylation changes.

First, let us set some notation. Methylation data associated to an individual, $j$, can be interpreted as a highly dimensional vector,

$$m_j = \left(m_j^0, \dots, m_j^S, t_j\right),$$

where $m_j^i$ are the methylation levels of individual $j$ at site $i$, and $t_j$ is the age of the individual. We denote by $\mathcal{D} = \{m_j\}_{j \in J}$ the collection of all data points associated to a cohort of individuals. We further assume that all data points $m_j^i$ are independently distributed both across individuals and sites. Finally, we consider the cross-section of all data points $\mathcal{D}^i = \left\{(m_j^i, t_j)\right\}_{j \in J}$, that is the data associated to a single CpG site for all individuals in the cohort.

We develop two mathematical models of methylation dynamics in a single CpG site and derive the predicted evolution in time of their associated probability distributions. Both models are fitted and compared using Generation Scotland's (GS) data. Then we derive the notion of acceleration and bias as the optimal perturbation at an individual level of the parameters inferred from GS.

#### 1 Biological model

To characterise the time-evolution of methylation patterns over time, mechanistically driven mathematical models offer a tangible interpretation of the underlying biological processes that drive these changes. At a single cell level, methylation changes in each genomic locus can be interpreted as a transition between unmethylated and methylated states, with rates $\nu_0$ and $\nu_1$ respectively (Fig. 2b),

$$U \to M, \quad \nu_0 \tag{1}$$
$$M \to U, \quad \nu_1 \tag{2}$$

This is an example of a simple continuous-time 2-state Markov chain that was first proposed in 1990 to study the clonal inheritance of CpG patterns [14], and more recently in the field of ageing in [17].

Let us denote by $X(t)$ and $Y(t)$ the number of unmethylated and methylated cells in the system at time $t$. The probability distribution of methylation levels in this system is given by

$$P\left(X\left(t\right)=a, Y\left(t\right)=b \mid X\left(0\right)=\left(1-p\right)N, Y\left(0\right)=pN\right), \tag{3}$$

where $N$ denotes the total number of cells in the system. Notice that we have split the initial population according to the proportion, $p$, of initially methylated cells $X\left(0\right)+Y\left(0\right)=\left(1-p\right)N+pN=N$. Further, since methylation array data only provides information about the proportion of methylated cells in the system, $m(t)=Y(t)/(X(t)+Y(t))$, and since the total number of cells in the system $X(t)+Y(t)$ is constant, we focus on inferring the distribution of the random variable $m(t)=Y(t)/N$.

Notice that, in general, there is no closed-form expression for the distribution of continuous-time Markov chains. We can however approximate the evolution of methylation patterns using a reparametrisation of the beta distribution in terms of a mean and variance, instead of $\alpha$ and $\beta$. More precisely, we model the probability distribution of $m(t)$ by

$$Beta\left(\mu_m(t), \sigma_m^2(t)\right),$$

where $\mu_m(t)$ and $\sigma_m^2(t)$ are mean and variance of $m(t)$. We can therefore, compute the probability of observing cohort data $\mathcal{D}^i$ in a single site, $i$, conditional on the proposed model of biological dynamics, as

$$P\left(\mathcal{D}^i|bio\right)=\prod_{j\in J} Beta_{pdf}\left(m_j^i, \mu_m(t), \sigma_m(t)\right).$$

In what follows we extract the moments of $X(t)$ and $Y(t)$ through the study of cumulant generating functions (cgf). To find the differential equation describing the cgf associated to the evolution of these random variables we lean on the technique described in Bailey ([3], Section 7.4). First, we write down the transition rates associated with the acquisition of $n$ and $m$ cells in each state $f_{n,m}$,

$$\begin{aligned} f_{-1,1} &= \nu_0 X(t) \\ f_{1,-1} &= \nu_1 Y(t) \end{aligned}$$

This yields the following differential equation for the cumulant generating function

$$\frac{\partial K}{\partial t}\left(\theta_0,\theta_1,t\right)=\frac{\partial K}{\partial \theta_0}\left(\theta_0,\theta_1,t\right)\nu_0\left(e^{-\theta_0+\theta_1}-1\right)+\frac{\partial K}{\partial \theta_1}\left(\theta_0,\theta_1,t\right)\nu_1\left(e^{\theta_0-\theta_1}-1\right)$$

with initial conditions

$$K\left(\theta_0,\theta_1,0\right)=pN\theta_0+\left(1-p\right)N\theta_1+c\theta_0^2+c\theta_1^2-c\theta_0\theta_1, \tag{4}$$

where $c$ denotes the variance of $X(t)$ and $Y(t)$ at time $t=0$. We can then easily obtain the differential equations for the cumulants of 3, taking derivatives of 1. Notice that taking derivatives with respect to $\theta_0$ and $\theta_1$ and valuing the resulting function at $\theta_0=\theta_1=0$, the left hand side of (1) becomes

$$\frac{\partial^{i+j}}{\partial\theta_0^i\partial\theta_1^j}\frac{\partial}{\partial t}K\left(0,0,t\right)=\frac{\partial}{\partial t}\left[k_{i,j}\left(t\right)\right].$$

More generally, we introduce the notation

$$f^{(i,j)}\left(\theta_0,\theta_1,t\right)=\frac{\partial^i}{\partial\theta_0^i}\frac{\partial^j}{\partial\theta_1^j}f\left(\theta_0,\theta_1,t\right),$$

and note that

$$\frac{\partial}{\partial t}K^{(i,j)}|_{\substack{\theta_0=0\\\theta_1=0}}=\frac{\partial k_{i,j}}{\partial t}\left(t\right),\forall i,j\geq 1,$$

where $\{k_{i,j}\left(t\right)\}$ corrrespond to the cumulants of the stochastic process. Further differentiating the right hand side of (1) and setting $\theta_0=\theta_1=0$ yields the differential equation satisfied by each cumulant.

#### 1.1 Expectation

To derive the time evolution of the expected evolution $\mathbb{E}[X(t)]$ and $\mathbb{E}[Y(t)]$ we consider the first order derivatives of the cumulants. It follows from the theory of cumulants that, in the multivariate case, $\mathbb{E}[X(t)]=k_{1,0}\left(t\right)$ and $\mathbb{E}[X(t)]=k_{0,1}\left(t\right)$, where $k_{i,j}(t)$ are defined through the differential equation (1). This yields the following differential equation for the expectation of $X(t)$ and $Y(t)$,

$$\begin{cases}\frac{d}{dt}k_{1,0}\left(t\right)= & -\nu_0k_{1,0}\left(t\right)+\nu_1k_{0,1}\left(t\right)\\\frac{d}{dt}k_{0,1}\left(t\right)= & \nu_0k_{1,0}\left(t\right)-\nu_1k_{0,1}\left(t\right)\end{cases}$$

with initial conditions $k_{1,0}\left(t\right)\left(0\right)=\left(1-p\right)N$ and $k_{0,1}\left(t\right)=pN$.

The analytical solution to this set of differential equations can be easily obtained using standard methods for solving ODEs and yields

$$\mathbb{E}[Y(t)]=N\eta_0+Ne^{-\omega t}\left\{\eta_0\left(p-1\right)+\eta_1p\right\},$$

where $\omega=\nu_0+\nu_1$ corresponds to the total rate of reactions in the system and $\eta_i=\nu_i/\omega$ to the proportion of methylation changes in each direction. Further, using the standard scaling of random variables and simplifying the above equation,

$$\mu_m(t)=\eta+e^{-\omega t}\left(p-\eta\right),$$

with $\eta = \eta_0$. Notice that the methylation dynamics $\mu_m(t)$ transition form an initial state,

$$\mu_m(0) = p,$$

towards a steady state that is independent of the initial state,

$$\lim_{t\to\infty} \mu_m(t) = \eta,$$

at a rate $\omega$. In other words, whilst the directionality of the process of ageing at a single site can be described by the initial and final state $p$ and $\eta$, the speed of ageing is given by the rate of transitions $\omega$.

#### 1.2 Variance

We now derive the analytical formula for the second order cumulants $k_{2,0}$, $k_{0,2}$ and $k_{1,1}$ corresponding to the variance of each population of cells, $X(t)$ and $Y(t)$ and their covariance.

Let us rewrite (1) as

$$\frac{\partial K}{\partial t}(\theta_0, \theta_1, t) = p(\theta_0, \theta_1)\frac{\partial K}{\partial \theta_0}(\theta_0, \theta_1, t) + q(\theta_0, \theta_1)\frac{\partial K}{\partial \theta_1}(\theta_0, \theta_1, t).$$

Using Leibniz rule, notation $p_0^{(i,j)} = p^{(i,j)}(0,0)$, and that $p(0,0) = q(0,0) = 0$, we find that

$$\begin{aligned}\frac{\partial k_{2,0}}{\partial t}(t) =& p_0^{(2,0)} k_{1,0}(t) + q_0^{(2,0)} k_{0,1}(t) \\ &+ 2p_0^{(1,0)} k_{2,0}(t) + 2q_0^{(1,0)} k_{1,1}(t),\end{aligned}$$

$$\begin{aligned}\frac{\partial k_{2,0}}{\partial t}(t) =& p_0^{(2,0)} k_{1,0}(t) + q_0^{(2,0)} k_{0,1}(t) \\ &+ 2p_0^{(1,0)} k_{2,0}(t) + 2q_0^{(1,0)} k_{1,1}(t),\end{aligned}$$

and

$$\begin{aligned}\frac{\partial k_{1,1}}{\partial t}(t) = & p_0^{(1,1)} k_{1,0}(t) + p_0^{(0,1)} k_{2,0}(t) + p_0^{(1,0)} k_{1,1}(t) \\ &+ q_0^{(1,1)} k_{0,1}(t) + q_0^{(0,1)} k_{1,1}(t) + q_0^{(1,0)} k_{0,2}(t),\end{aligned}$$

We will simply give the expression of the relevant $p_0^{(i,j)}$ and $q_0^{(i,j)}$ and urge the reader to substitute the formulas to obtain the explicit expression of the differential equations associated to the second order cumulants. Recall the expressions for $p$ and $q$

$$p(\theta_0, \theta_1) = \nu_0\left(e^{-\theta_0+\theta_1} - 1\right)$$

and

$$q(\theta_0, \theta_1) = \nu_1\left(e^{\theta_0-\theta_1} - 1\right)$$

Then

$$
\begin{aligned}
p_0^{(1,0)} =& -\nu_0 \\
p_0^{(0,1)} =& \nu_0 \\
p_0^{(2,0)} =& \nu_0 \\
p_0^{(1,1)} =& -\nu_0 \\
p_0^{(0,2)} =& \nu_0
\end{aligned}
$$

and using the symmetry of $p$ and $q$, it easily follows that

$$
\begin{aligned}
q_0^{(1,0)} =& \nu_1 \\
q_0^{(0,1)} =& -\nu_1 \\
q_0^{(2,0)} =& \nu_1 \\
q_0^{(1,1)} =& -\nu_1 \\
q_0^{(0,2)} =& \nu_1
\end{aligned}
$$

Using the initial conditions, (4),

$$
\frac{\partial k_{2,0}}{\partial t}(0) = \frac{\partial k_{0,2}}{\partial t}(0) = c
$$

and

$$
\frac{\partial k_{1,1}}{\partial t}(0) = -c,
$$

we can solve the initial boundary problems using standard methods for coupled ODEs.

We finally obtain the time evolution of the variance of the proportion of methylated cells $m(t)$,

$$
\sigma_m^2(t) = \frac{\eta_0\eta_1}{N} + e^{-\omega t}\left\{\frac{\eta_0^2(1-p) + p\eta_1^2 - \eta_0\eta_1}{N}\right\} + e^{-2\omega t}\left\{\frac{c}{N^2} + - \frac{\eta_0^2(1-p) + p\eta_1^2}{N}\right\}.
$$

Notice that as expected, the initial variance of the system is given by

$$
\sigma_m^2(0) = \frac{c}{N^2},
$$

and evolves towards a limiting state

$$
\lim_{t\to\infty} \sigma_m^2(t) = \frac{\eta_0\eta_1}{N},
$$

#### 1.3 System size

Notice that the system size $N$ does not play any role in the evolution of the mean of $m(t)$ and simply plays a scaling role in the evolution of $\sigma^2(t)$. That is, the system size only controls the amount of stochastic variance in the modelled system, where higher values of $N$ will result in a lower variance and vice-versa. Since our model is a simplification of the true process of methylation changes, we are likely underestimating the true system size, to artificially increase the level of complexity in the system (or stochastic variance).

##### 1.4 First order Taylor expansion

Notice that the proposed biological model predicts an exponential evolution of both the mean and variance of $m(t)$. However, linear clocks have successfully recapitulated to a great degree the evolution of methylation levels throughout the lifespan of an individual. The discrepancy between this model's dynamics and those predicted by linear models can be explained as CpG sites included in current epigenetic predictors, will have have a small rate of reactions $\omega \ll 1$. In that regime, we can use the first order Taylor expansion of the exponential

$$e^{-}\omega t \sim (1 - \omega t)$$

to develop a linear approximation to the behaviour of methylation patterns in a linear model. In that case,

$$m(t) \sim p - \omega t(p - \eta),$$

shows an initial behaviour close to the biological model, with initial state $m(0) = p$ , that diverges from the model as $t$ increases.

Notice however, that In this model it already becomes clear that an increase in speed of methylation, $\tilde{\omega} = \alpha\omega$ changes is associated with an increase of the slope of the linear approximation.

Similarly, the evolution of the variance can be approximated and, in the regime where $e^{-2\omega t} \sim (1 - 2\omega t)$, we have

$$\sigma_m^2(t) = \frac{\eta_0\eta_1}{N} + (1 - \omega t) \left\{ \frac{\eta_0^2(1-p) + p\eta_1^2 - \eta_0\eta_1}{N} \right\} + (1 - \omega t)^2 \left\{ \frac{c}{N^2} - \frac{\eta_0^2(1-p) + p\eta_1^2}{N} \right\}$$

This suggest that linear modelling of methylation changes should at least contain linearly increasing variance term.

#### 2 Linear model

As suggested by the Taylor approximation of our biological model and previous epigenetic clocks, the distribution of methylation values at age $t$ can be approximated by a normal distribution with a mean that increases linearly with time and a constant variance. This is indeed the model used in widespread predictors such as Hannum's or Horvath's epigenetic clocks , [8] and [9] respectively. In these models, the probability of observing a methylation value in a single CpG site $i$ at age $t$ is given by the random variable $M^i(t)$,

$$M^i(t) \sim \mathcal{N}\left(a^i t + b^i,\ c^i\right),$$

where $a_i, b_i$ and $c_i$ are the model parameters.

On a cohort level the probability of observing cohort data $\mathcal{D}^i$ in a single site, $i$, conditional on model of linear dynamics, as

$$P\left(\mathcal{D}^i | bio\right) = \prod_{j \in J} Beta_{pdf}\left(m^i_j, \mu_m(t), \sigma_m(t)\right).$$

#### 2.1 Limitations - Global bias

Note that in a linear model, predictions of epigenetic age are given by

$$\hat{y} = \sum_{i \in I} a^i m^i_j t_j + b^i,$$

where $m^i_j$ are the methylation levels of individual $j$ in sites $i \in I$, and $t_j$ the age of the individual. Current epigenetic predictors then compute the epigenetic acceleration as the distance from the prediction to chronological age, therefore

$$\begin{aligned} acc_j &= \hat{y} - t_j && (5) \\ &= -t_j + \sum_{i \in I} a^i m^i_j t_j + b^i && (6) \end{aligned}$$

We no consider the effect of a global change in methylation on our acceleration prediction. To simplify the interpretation we focus on an individual whose prediction is perfect, that is 0 acceleration. For this individual,

$$t_j = \sum_{i \in I} a^i m^i t_j + b^i.$$

We then modify $m^i_j$ by a global offset independent of the site $i$,

$$\tilde{m^i_j} = m^i_j + \beta_j.$$

The prediction acceleration for this shifted individual then becomes

$$\begin{aligned} a\tilde{c}c_j &= \hat{y} - t_j && (7) \\ &= -t_j + \sum_{i \in I} a^i \tilde{m^i_j} t_j + b^i && (8) \\ &= -t_j + \left(\sum_{i \in I} a^i m^i_j t_j + b^i\right) + \left(\sum_{i \in I} a^i \beta_j t_j\right) && (9) \\ &= \beta_j \sum_{i \in I} a^i t_j && (10) \end{aligned}$$

More generally the acceleration predictions are shifted as follows

$$a\tilde{c}c_j = acc_j + \beta_j \sum_{i \in I} a^i t_j.$$

### 3 Model comparison

We use the Generation-Scotland's cohort data to approximate the posterior distribution of both the biological and linear model using Markov Chain Monte-Carlo algorithms implemented in PyMC [16]. Further, we can compare both models using the approximated the expected log-predictive density (ELPD) of both models on each site using an approximation of leave-one-out cross validation (LOO-CV) based on the Pareto-smoothed importance sampling (PSIS).

### 4 Modelling acceleration and bias

To compute the acceleration and bias for each individual in a cohort we first extract he maximum likelihood estimator parameters for the biological model, $\overline{\omega_i}, \overline{\eta_i}, \overline{N_i}, \overline{p_i}$ and $\overline{c_i}$ associated to each CpG site.

That is, notice that in (1.1) and (**??**),

$$\begin{aligned} \mu_m(t) &= \eta + e^{-\omega t}(p-\eta) && (11)\\ &= \mu_m(t,\omega,p,\eta) && (12) \end{aligned}$$

and similarly, $\sigma^2(t) = \sigma^2(t,\omega,\eta,p,N,c)$. Then we use cohort data $\mathcal{D}^i$ to compute the maximum likelihood estimators $\overline{\omega_i}, \overline{\eta_i}, \overline{N_i}, \overline{p_i}$ satisfying

$$P\left(\mathcal{D}^i|\overline{\omega_i},\overline{\eta_i},\overline{N_i},\overline{p_i},c_i\right) = \max_{\omega^i,\eta^i,p^i,N^i,c^i}\prod_{j\in J} Beta_{pdf}\left(m_j^i,\mu_m(t_j,\omega^i,p^i,\eta^i),\sigma^2(t_j,\omega^i,\eta^i,p^i,N^i,c^i)\right).$$

We simplify the notation by setting $\theta^i = \left\{\omega^i,\eta^i,p^i,N^i,c^i\right\}$ and define $\overline{\theta^i}$ similarly.

Then, given an in individual $j$ and its associated data

$$m_j = \left(m_j^0,\ldots,m_j^S,t_j\right),$$

where $m_j^i$ corresponds to the methylation levels at site $i$ and $t_j$ the age of the individual, we can compute the probability of observing its methylation pattern

$$P\left(m_j\right) = \prod_{i=0}^{S} Beta_{pdf}\left(m_j^i,\mu_m(t,\overline{\theta^i}),\sigma_m^2(t,\overline{\theta^i})\right),$$

We can then consider the effect of increasing the speed of methylation changes uniformly across CpG sites on this probability. That is, to consider the probability associated with

$$\overline{\theta^i}(\alpha,\beta) = \left\{\overline{\omega^i}(\alpha),\overline{\eta^i}(\beta),\overline{p^i}(\beta),N^i,c^i\right\}$$

where $\overline{\omega^i}(\alpha) = \alpha * \overline{\omega^i}$ denotes a proportional increase in the total rate of transitions and $\overline{p^i}(\beta) = \overline{p^i} + \beta$ and $\overline{\eta^i}(\beta) = \overline{\eta^i} + \beta$ global shifts in the initial and final states. Notice that the resulting average evolution of methylation changes in site $i$ is given by

$$\mu_m\left(t, \overline{\theta^i}(\alpha,\beta)\right) = \overline{\eta^i} + \beta + e^{-\alpha\overline{\omega^i}t}\left(\overline{p} - \overline{\eta}\right).$$

We can then infer the acceleration $\alpha_j$ and bias $\beta_j$ for every individual as the maximum likelihood estimators satisfying

$$P\left(m_j, \alpha_j, \beta_j\right) = \max_{\alpha,\beta} \prod_{i=0}^{S} Beta_{pdf}\left(m_j^i, \mu_m(t, \overline{\theta^i}(\alpha,\beta)), \sigma_m^2(t, \overline{\theta^i}(\alpha,\beta))\right).$$

### 5 Batch correction

We now proceed to define an algorithm of batch correction that allows to transfer the optimal model parameters $\overline{\theta^i}$ inferred using GS for each site to other cohorts. To do so, we simply assume that across cohorts, the dynamics in CpG sites remain largely unaltered but show a shift in the mean that is independent for each site. That is, for a new set of data $\tilde{\mathcal{D}}^i$ associated to a CpG site $i$, we can compute

$$P\left(\tilde{\mathcal{D}}^i|\Omega^i\right) = \prod_{j\in J} Beta_{pdf}\left(m_j^i, \mu_m(t, \overline{\theta^i}) + \Omega^i, \sigma_m^2(t, \overline{\theta^i})\right),$$

where $\Omega^i$ denotes the global offset in site $i$. We then infer the optimal value maximizing the likelihood of $\Omega^i$, to shift the predicted dynamics in the new cohort.

### 6 Quality control of Generation Scotland data

#### 6.1 DNA methylation data

Whole blood genomic DNA samples were normalised to 50 nanograms per microlitre and were treated with sodium bisulfite using the EZ-96 DNA Methylation Kit (Zymo Research, Irvine, California), following the manufacturer's instructions. DNA methylation was profiled using the Infinium MethylationEPIC BeadChip. Methylation typing was performed in 31 batches within each set.

In Set 2, the R ShinyMethyl package was used to compare plots of log median signal intensities across methylated and unmethylated beads in each array [7]. Outliers were removed based on a visual inspection of the plots. The wateRmelon package in R was used to remove samples based on the following exclusion criteria: (i) >1% of probes had a detection p-value > 0.05, (ii) probes had a beadcount 5% of samples and (iii) probes were non-autosomal [15]. Eighty samples and 5,910 probes were excluded based on these criteria. Probes that were predicted to have off-target effects were excluded along with those that contained a SNP in the final five 3' bases or at the site of single-base extension in type I probes (n = 84,352) [11, 19]. Individuals were removed if (i) their methylation-based predicted sex did not match recorded sex (n=12), (ii) their sample was derived from salive (n=10), (iii) they were genetic outliers in PCA of genotype data (n=7) or (iv) they self-reported 'Yes' to all health conditions listed on study questionnaires at baseline (n=3). One individual was also removed as their methylation data indicated that they might have an XXY genotype. Data were normalised using the dasen method in the wateRmelon package. The final set consisted of 5,087 participants and 760,943 loci [18]. In Set 2 there were 2,578 unrelated individuals brought forward for analyses in this study.

In Set 1, the Meffil package in R was used to perform initial quality control steps [12]. Samples were excluded if they met the following criteria: (i) there was a mismatch between self-reported and methylation-based predicted sex, (ii) more than 1% of probes had a detection p-value$>$ 0.05, (iii) samples showed evidence of dye bias, (iv) sample were outliers at bisulfite conversion control probes and (v) the sample had a median methylated signal intensity that was $\geq$ 3 standard deviations lower than expected. ShinyMethyl was then used to exclude probes based on the same criteria as those applied in Set 2. Meffil was re-employed to remove poor-performing probes that met the following criteria: (i) probes had a beadcount of 5% of samples and (ii) $>$ 5% of samples had a detection p-value $>$ 0.05. In total, 8,878 poor-performing probes and 135 samples were excluded. Probes with potential off-target effects were excluded as in Set 1, again along with those that had a SNP in the final five 3' bases or in the single-base extension site (n = 84,352). Data were normalised using the dasen method. There were 4,450 individuals in Set 1 and 758,332 CpG sites following quality control procedures. The individuals in Set 1 were unrelated to each other and to those in Set 2.

#### 6.2 Genotype data

GS genotyping was carried out over two batches with 9863 samples genotyped using the Illumina HumanOmniExpressExome-8 v1.0 Bead Chip with the remainder genotyped using the Illumina HumanOmniExpressExome-8 v1.2 Bead Chip. Infinium chemistry was employed in both batches. Quality control was carried out in PLINK v1.9b2c [4]. SNPs with a call rate $<$ 0.98, minor allele frequency $\leq$ 0.01 and Hardy-Weinberg equilibrium test with $p \leq 10^{-6}$ were removed. There was a total of 561,125 autosomal SNPs that passed quality control. Duplicate samples were removed. Samples were also excluded if they had a genotype call rate $<$ 0.98. PCA of genotype data was performed to identify potential outliers. GS genotypes were combined with data from 1,092 individuals in the 1000 Genomes Project before PCA [6]. Outliers who were more than six standard deviations away from the mean of the first two principal components were removed [2]. Following quality control there were 19,904 individuals with genotype data, consisting of 11,731 females and 8,173 males [10, 5]. Genotype data were imputed using the Haplotype Reference Consortium panel. Monogenic and low imputation quality (INFO$<$ 0.4) variants were removed from the imputed dataset leaving 24,161,581 variants for downstream analyses [13, 1].
